# Overexpression of Rheb, a positive Tor regulator, reveals principles of rod outer segment size control

**DOI:** 10.1101/2023.09.08.556898

**Authors:** Larissa Gilliam, John J Willoughby, William Annan, Diana White, Abbie M Jensen

## Abstract

The outer segment of retinal rod photoreceptors is a highly modified cilium with the typical ciliary microtubular axoneme but additionally there are hundreds of densely packed, stacked, discrete intramembranous discs containing phototransduction machinery. The outer segment is continuously regenerated or renewed throughout the life of the animal through the combined processes of proximal growth and distal shedding. Since the pioneering experiments decades ago that demonstrated the process of rod outer segment renewal very little research has been directed towards understanding mechanistic principles of renewal and homeostasis, and many important questions remain unaddressed. The rod outer segment is a prime example of tight size control that is amenable to experimental and computational analyses. This study uses zebrafish molecular genetics with high resolution imaging, quantitative analyses, and computational modeling to uncover principles that regulate rod outer segment homeostatic length control. By genetically increasing rod outer segment growth, we show that outer segments lengthen and shedding does not accelerate to offset this added growth.

## Introduction

The mechanisms used by cells to measure and control organelle size remain poorly understood. The outer segment of rod photoreceptors is a fascinating and tractable example of tight, continual size control. Vertebrate photoreceptors are highly compartmentalized postmitotic neurons that consist of four functional compartments: two basal compartments– the synaptic region and the cell body, and two apical compartments– the inner segment and the outer segment. The rod outer segment (ROS) is a highly modified primary cilium where photons of light are captured by Rhodopsin proteins and transduced into changes in membrane potential that alter synaptic neurotransmitter release. The ROS has the typical ciliary microtubular axoneme but additionally holds hundreds of densely packed, stacked, discrete intramembranous discs containing the phototransduction machinery (1300-1500 discs in a human ROS) enclosed by the plasma membrane [1]. The ROS is about 10 times longer, volume about 250-300 times greater, and membrane area nearly 1000 times greater than a typical primary cilium [2] [3]. Another major difference between photoreceptor outer segments and primary cilia is that the outer segment is continuously regenerated, or renewed, throughout the life of the animal through the combined daily processes of proximal growth and distal shedding, the retinal pigmented epithelium (RPE) phagocytoses and digests the ROS tips [4] [5] [6] [7] [8]. It is postulated that photoreceptors renew their outer segments because the very narrow connecting cilium cannot accommodate the retrieval of worn outer segment disc membrane and associated proteins for disposal and recycling in the inner segment.

Since the pioneering experiments decades ago revealing the process of ROS renewal very little research has been directed towards understanding mechanistic principles of outer segment renewal and length control, and many important questions remain unaddressed. What is the growth machinery that adds new material at the ROS base? What is the shedding machinery at the ROS tip? What determines growth and shedding rates? Is there an outer segment length set-point? If so, how is it set and what molecules determine the set-point? Are growth and shedding coordinately regulated? Using a transgenic zebrafish line that uniquely allows quick and quantitative measures of both ROS growth and shedding rates together with doxycycline (DOX)-induced transgene expression, we designed and performed experiments to uncover principles of ROS renewal and size control. We sought to increase ROS growth by overexpressing *rheb*, a positive regulator of the Tor pathway of cell size control, in rods [9].

Here we report that ROS growth increases upon Rheb-overexpression, but shedding does not increase to offset the added growth, suggesting that a simple set-point model of length control is insufficient. Additionally, we find the ROS axoneme does not lengthen concomitantly with ROS length elongation, suggesting that axoneme length does not directly control ROS length. Additionally, to complement our experimental observations, and to provide further insight into the mechanism(s) of ROS length control, we develop a mathematical model of the ROS renewal process. By varying parameters in our computational model that increase the growth dynamics of the ROS (i.e., similar to genetic Rheb overexpression), results show statistically significant similarities between experimental observations and simulation results.

## Results

To measure ROS renewal dynamics in this study, we use the *Tg(hsp70:HA-mCherry^TM^)* transgenic line in which a brief increase in water temperature induces expression of transmembrane mCherry protein that is incorporated into a small number of newly synthesized discs at the base of the ROS, these labeled discs are displaced distally as new, unlabeled discs are added [10]. This method allows us to quantify growth rates and shedding rates (Fig. 1A); the red discs represent mCherry labeling at the time of heat-shock induction. The distance from the base of the ROS to the stripe is growth post heat-shock, whereas the distance from the stripe to the tip is growth before heat-shock minus subsequent shedding.

**Figure 1.**
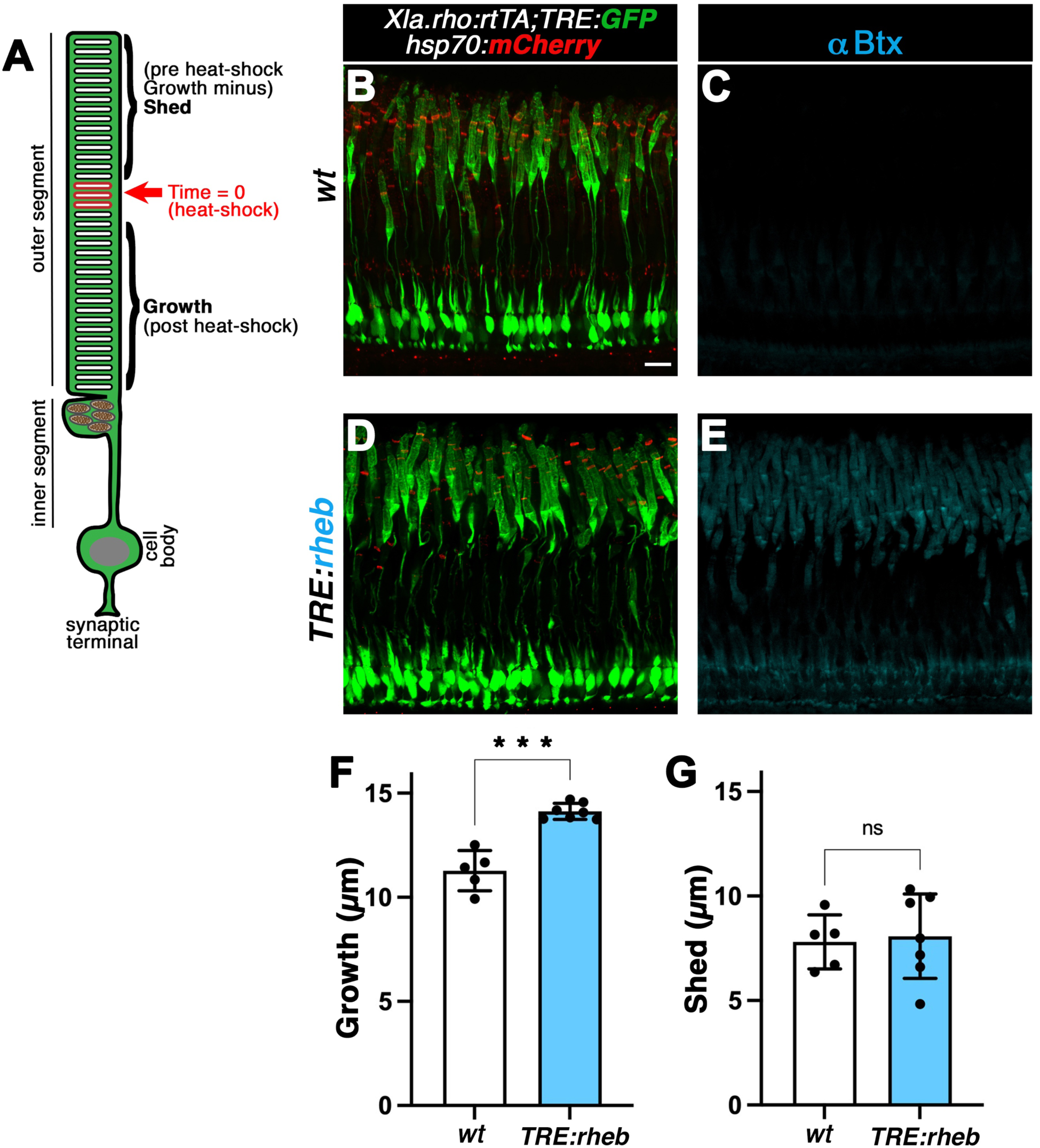
Rheb-overexpression increases ROS growth but does not affect ROS shedding. **(A)** Diagram of the rod photoreceptor structure illustrating the outer segment– a highly modified primary cilium with stacked discrete intramembranous discs, the inner segment with a unique population of giant mitochondria concentrated at the base of the outer segment, the cell body and the synaptic terminal. The red discs represent mCherry labeling by heat-shock induction. The distance from the base of the outer segment to the stripe is growth post heat-shock. The outer segment from the stripe to the tip is outer segment length that was present before heat-shock minus subsequent shedding. **(B, D)** Confocal z-projections (z = 4 μm) of the photoreceptor layer in *wild-type* (*wt*) (**B**) and *TRE:rheb* (**C**) transgenic retinas showing GFP expression in rods and heat-shock induced mCherry stripe after 14 days of DOX treatment. Scale bar, 10 μm. **(D, E)** Confocal z-projections (z= 4 μm) of untreated *wild-type* (*wt*) (**D**) and *TRE:rheb* transgenic (**E**) retinas, labeled with αbungarotoxin (αBtx) and ROS labeled with anti-Rhodopsin (anti-Rho) antibodies. Nuclei labeled with DAPI. Scale bar, 10 μm. (**F**) Quantification of growth in *wild-type* (*wt*) and *TRE:rheb* transgenic ROS, after 14 days DOX treatment. Growth was measured as the distance from the base of the ROS to the red stripe. Each point is the average of measurements from an individual eye. Growth is significantly (*p* < 0.0016) greater in *TRE:Rheb* transgenic ROS. Error bars, SD; Significance, Welch’s *t*-test. (**E**) Quantification of shedding in *wild-type* (*wt*) and *TRE:rheb* transgenic ROS, after 14 days DOX treatment. Shedding was measured as the distance from the red stripe to the ROS tip. Each point is the average of measurements from an individual eye (error bars, SD; Significance, Welch’s *t*-test).

To temporally control *rheb* expression exclusively in rods, we used the rod Tet-On driver line, *Tg(Xla.rho:rtTA;TRE:GFP)* and created the *Tg(TRE:rheb^aBtx^)* transgenic response line, and combined them with the *Tg(hsp70:HA-mCherry^TM^)* transgenic line [10] [11]. Fish were in the *albino* background as dense RPE microvillar melanin heavily obscures the photoreceptor outer segment region. Adult *Tg(Xla.rho:rtTA;TRE:GFP); Tg(hsp70:HA-mCherry^TM^); alb^-/-^* with and without *Tg(TRE:rheb^aBtx^)* were heat-shocked and immediately treated with doxycycline (DOX) for 14 days. We acquired confocal z-stack images of rod photoreceptors (Fig. 1B, D) and measured growth rates (distance from the ROS base to the mCherry stripe) and shedding rates (distance from the mCherry stripe to the ROS tip; Fig. 1A). To reveal Rheb over-expression, we labeled the tissue with αBtx (Fig. 1 C, E). We also labeled tissue with αBtx in DOX-untreated *Tg(Xla.rho:rtTA;TRE:GFP); Tg(hsp70:HA-mCherry^TM^); alb^-/-^; Tg(TRE:rheb^aBtx^)* fish, which shows no Rheb transgene expression without DOX treatment (Supplemental Fig 1). We find that Rheb-overexpression increases ROS growth by approximately 25% (Fig. 1F), while shedding is not significantly different from *wt* rods (Fig 1G).

We next examined whether the microtubule-based axoneme, which extends partway along the ROS and is a good candidate to regulate ROS length (Fig 2A), is affected by Rheb-overexpression. We acquired confocal z-stack images of anti-acetylated Tubulin labeled retinal tissue and measured axoneme length in animals heat-shocked and DOX-treated for 14 days (Fig 2B-E). We observe no significant difference in axoneme length between Rheb-overexpressing rods and *wt* rods (Fig 2F).

**Figure 2.**
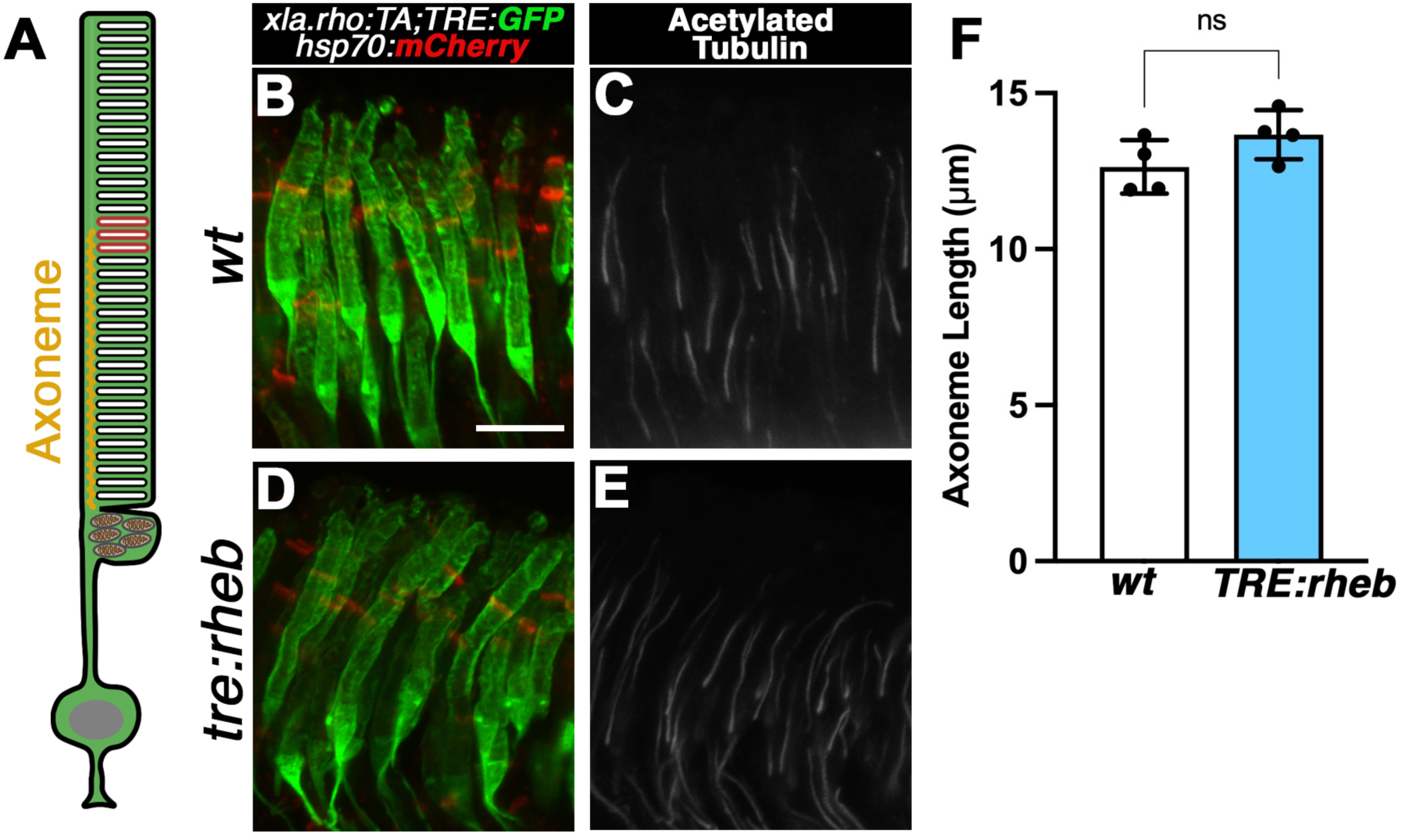
ROS ciliary axoneme length is unaffected by Rheb overexpression and increased ROS growth. (**A**) Diagram of rod photoreceptor illustrating the ciliary axoneme (yellow) extending partway through the outer segment. (**B-E**) Confocal z-projections (z = 4 μm) of GFP expression, heat-shock induced mCherry stripe and anti-acetylated Tubulin labeling after 14 days of DOX-treatment in *wild-type* (*wt*) (**B**, **C**) and *TRE:rheb* (**D**, **E**) in ROS. Scale bar, 10 μm. (**F**) Quantification of axoneme length in *wt* and *TRE:rheb* ROS. Each point is the average of axoneme measurements from an individual’s eye. Error bars, SD; Significance, Welch’s *t*-test.

Loss of mTor negative regulators, Tsc1 and Tsc2, can lead to longer primary cilia, as measured by axonemal labeling [16] [17] [18] [19]. Our data show the ROS axoneme does not lengthen significantly upon Rheb-overexpression, thus, the ROS axoneme is a poor candidate to regulate ROS size control (Fig 3B).

**Figure 3.**
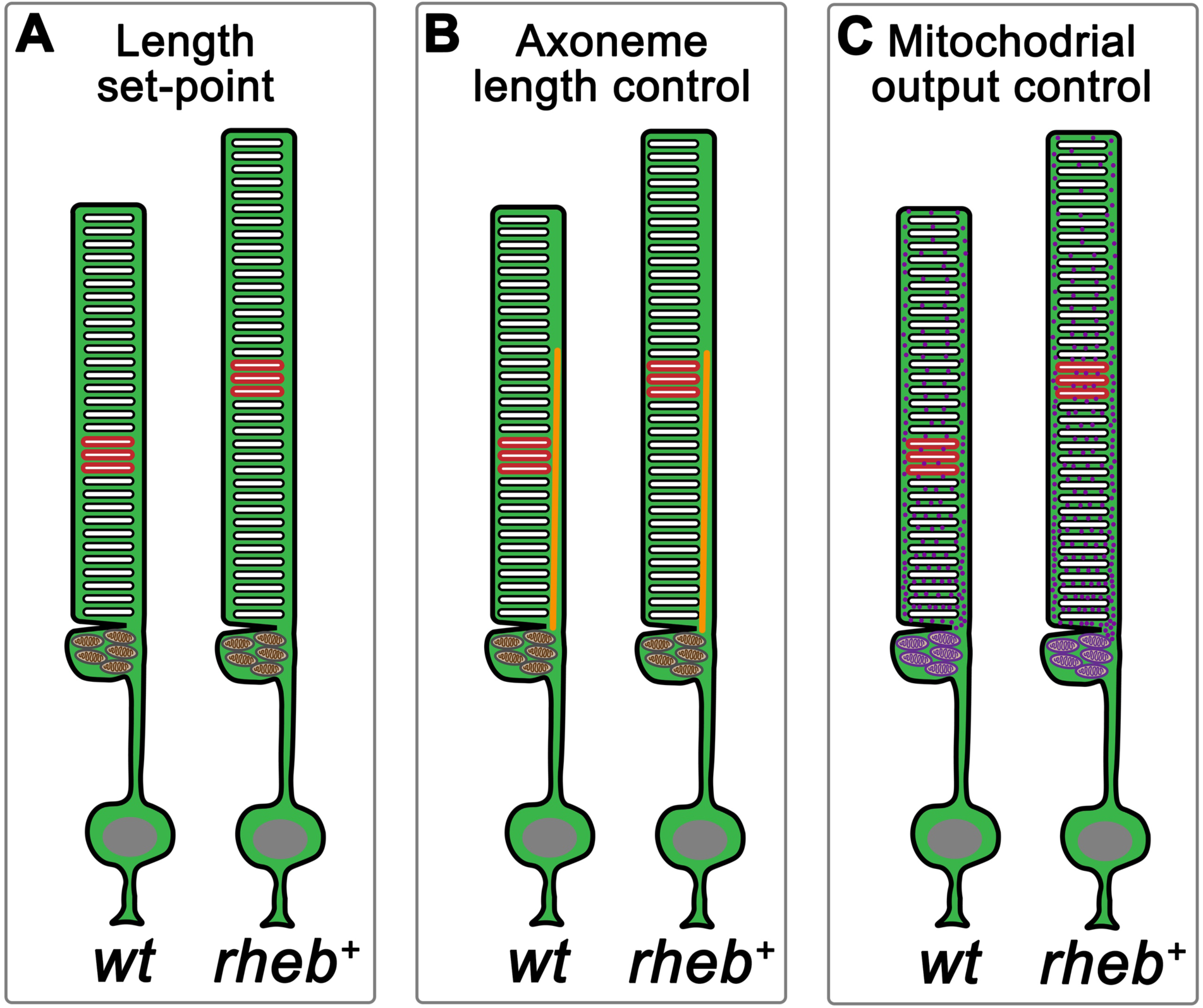
Models of ROS length control. **(A)** The set-point model of homeostatic ROS length control proposes that there exists a pre-determined length to which growth and shedding maintain. Rheb-overexpression results in ROS that are longer than *wt* ROS, thus the simple set-point model is rejected. **(B)** The ROS axoneme model of homeostatic ROS length control proposes axoneme length determines overall ROS length, perhaps by transporting cargo toward the ROS tip. Rheb-overexpression did not significantly increase axoneme length, thus, the axoneme model is rejected. **(C)** The mitochondrial function model of homeostatic ROS length control proposes that the ATP output from giant and abundant mitochondria at the base of the ROS regulates ROS length. The mitochondrial model is consistent with ROS elongation by Rheb-overexpression. Future experiments that enhance mitochondrial function can directly test this model.

Mitochondrial function is stimulated by mTor activity [20]. The ellipsoid region of the inner segment sits at the base of the outer segment and contains distinctive, numerous, giant mitochondria [21] [22]. These mitochondria are well positioned to metabolically support ROS function and regulate ROS length, by both increasing disc addition at the base and supporting integrity at the ROS tip (Fig 3C). A mitochondria-centered model of ROS length control is consistent with our data and also with the observations in the experiments described for the ROS ‘set-point’ length model described above. Further, the mitochondria-centered model of ROS length control is consistent with the observation that the ROS progressively shortens before photoreceptors die in mouse models of human retinal degeneration disease [23] [24] [25] [26] [27] [28] [29] [30].

To model ROS renewal kinetics, we formulated a mathematical model which consists of a system of three ordinary differential equations (ODEs), based on the schematic diagram in Figure 4A. We partition the system into three distinct compartments, the first being the growing compartment “G” located at the very base of the ROS. Discs in this compartment are still connected to the cell membrane, where they are open to the extracellular space and have not attained full disc diameter. We assume that new discs are added to the G compartment at a disc addition rate µ_g_ [31], which describes the average number of discs added to the ROS each day. The second compartment is the mature compartment “M” consisting of fully developed discs; those translocated toward the distal end of the ROS. We assume that the rate of maturation of new discs from the G to M compartment is governed by the rate of translocation of discs along the rod, the maturation (growth) rate α_g._ The discs are then shed from the M compartment to the shed compartment “S” at the shedding rate α_s_, being engulfed by the RPE. Finally, shed discs are disposed by the RPE at disposal rate µ_s_. Although there is no experimental work that describes the disposal rate, biologically this function exists. Our computational model requires the disposal term so that the number of discs being disposed of in the S compartment does not become too large.

**Figure 4.**
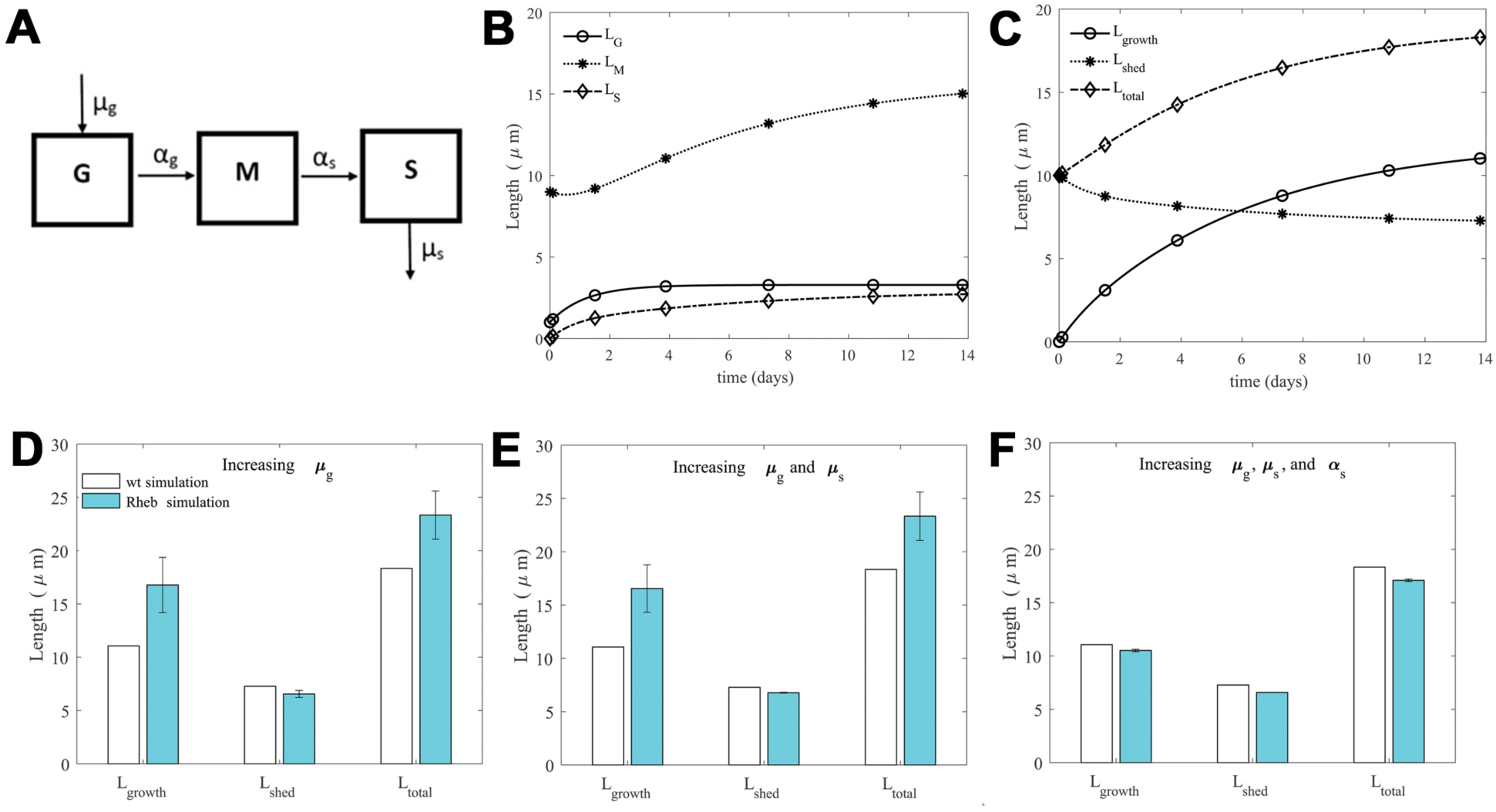
Model construction, model output, and comparison with experimental data. **(A)** A compartmental diagram showing the relationship and the flow of information between the three compartments used in building the model. G is the growing compartment (above mitochondria and at the base of ROS), M is the mature compartment, only composed of mature discs, and S is the shed compartment, where the RPE disposed of shed discs. Here, we define the parameters *µ_g_* as the disc addition rate, *α_g_* as the disc maturation rate*, α_s_* the shedding rate, and *µ_s_* the rate at which discs are disposed after being shed. **(B)** Model output for the time evolution of the three compartments (G, M & S) where we convert disc number to length using equation (1) (equations describing the time evolution of these three compartments (the ODEs) are found in the Supplementary Material). We simulate the model using the baseline parameter set: *µ_g_* = 82.0 discs/day, *α_g_* = 0.85 μm/day, *µ_s_* = 0.96 μm/day, and *α_s_* = 0.17 μm/day. This baseline parameter set, which corresponds to our simulated *wt* parameter set, is obtained from the parameter sweep described in the Supplementary Material. The initial condition for the model is G(0) = 30 discs, M(0) = 272 discs, and S(0) = 0 discs, which converted to lengths using equation (1) gives L_G_(0) = 1 μm, L_M_(0) = 9 μm, and L_S_(0) = 0 μm. **(C)** The corresponding lengths for L_growth_, Ls_hed_, and L_total_ over time from the results found in (B).| Disc to length conversion completed using equations (4), (3), and (2). At day 14, the model results in L_growth_ = 11.1 μm, L_shed_ = 7.2 μm, and L_total_ = 18.2 μm. **(D-F)** White bars are baseline values recorded for L_growth_, L_shed_, and L_total_ at day 14 (those values described in (C)). Blue bars are mean and SD of L_growth_, L_shed_, and L_total_ at day 14 for 50 runs of each simulation tested. For each simulation test, we varied model parameters to explore the mitochondrial hypothesis. **(D)** Increasing the disc addition rate, µ_g_, from its baseline value in the *rheb* simulation results in increased L_growth_ and L_total_. We completed 50 runs of the simulation, where we selected random values for µ_g_ from a uniform distribution with SD = ± 75 discs/day and mean higher (mean µ_g_ = 106 discs/day) than the baseline value (mean µ_g_ = 82 discs/day). **(E)** Increasing both the disc addition rate, µ_g_, and the disposal rate, µ_s_, from baseline values in the *rheb* simulation results in increased L_growth_ and L_total_ (similar to **D**). We completed 50 runs of the simulation, where for each run we selected random values for µ_g_ (as described in E), and µ_s_ from a uniform distribution with SD = ± 0.25 μm/day and mean higher (mean µ_s_ = 1.06 μm/day) than the baseline value (mean µ_s_ = 0.96 μm/day). **(F)** Increasing the disc addition rate, µ_g_, the disposal rate, µ_s_, and the shedding rate, α_s_, from baseline values in the *rheb* simulation results in small decreases in the growth and shed lengths. We completed 50 runs of the simulation, where for each run we selected random values for µ_g_ (as described in E), µ_s_ (as described in F), and α_s_ from a uniform distribution with SD = ± 0.65 μm/day and mean higher (mean α_s_ = 0.27 μm/day) than the baseline value (mean α_s_ = 0.17 μm/day).

Our model describes the rate of change of discs in each of the G, M, and S compartments (see Supplementary Materials for details). To illustrate our results, we show results in terms of *length*, as opposed to disc number, to better compare with experimental observations. The conversion of disc number to length is given by equation (1) and is based on experimental disc-disc spacing measurements [32] (see Materials and Methods). Figure 4B shows a simulation of the model, illustrating how length for each compartment changes over 14 days. Here, we run our model using what we call our “baseline” parameter set (described in detail below), which is meant to represent ROS dynamics in a *wt* setting. We stop our model simulation at 14 days, as this was the time that lengths were recorded in experiments following DOX-treatment. In our model, time t = 0 corresponds to the moment of heat-shock, where the initial length of the ROS is determined by adding the lengths associated with the G and M compartments at this time. Figure 4C illustrates how the shed length (which corresponds to the initial length minus that which is lost to the S compartment by subsequent shedding after heat shock), the growth length (which corresponds to the total length minus that shed), and the total length change over time. See Materials and Methods for equations (3) and (4) that describe how shed and growth lengths are calculated.

To test both the set-point hypothesis and the mitochondrial hypothesis, we varied model parameters for which we assumed would be influenced for each hypothesis (parameters varied in each simulation will be described below). Before varying model parameters, we determined a fixed parameter set, which we call our baseline parameter set, for which our model produces growth and shortening lengths similar to the *wt* experiments found in Figs. 1F and 1G, respectively, at 14 days. Here, we fixed the maturation rate α_g_ = 0.85 µm/day (which corresponds to the average growth length over 14 days from Fig. 1F), and completed a parameter sweep over a large range for each of the other three parameters, recording all sets of parameters that gave a growth and shed length within the experimental range at 14 days. Then, we selected the average of each parameter to be our fixed/baseline parameter set (see Supplementary Materials for more details on the ranges used for parameters in the parameter sweep and average values for each parameter, which we defined as our baseline parameter set).

All parameter variations, regardless of the test we were performing, were completed in a similar fashion. That is, we selected random values for each parameter in question, from a uniform random distribution, whose mean was higher than that of its baseline value. Figure 4D-F show the results (mean growth length, shed length, and total length; blue bars) for Rheb overexpression where each simulation was run 50 times up to 14 days, and the *wt* results (growth length, shed length, and total length; white bars) from the fixed parameter set at 14 days.

The set-point hypothesis is not supported by an examination of the model’s equilibrium, the values for G, M, and S that our model predicts as an eventual fixed solution (no longer varying in time, described in the Supplementary Materials). For our model, G, M, and S will approach the equilibrium values G = µ_g_/α_g_, M = µ_g_/α_s_, and S = µ_g_/µ_s_ discs as time progresses. Thus, if we increase the disc maturation rate α_g_, together with the disc shedding rate α_s_, the equilibrium result suggests a decrease in the G and M values (i.e., a decrease in the overall ROS length).

To test the mitochondrial hypothesis, we first examined how/if increases in the disc addition rate µ_g_ from baseline (keeping all other parameters fixed at baseline) would alter the growth and shed lengths at 14 days. Here, we suggest that an increase in disc addition rate at the base of the ROS might correlate with increased mitochondrial activity. In Figure 4D, we observe a significant increase in the growth length, *L_growth_*, about 40% greater than *wt* baseline (white bar), similar to the significant increase in *L_growth_* described by *rheb*-overexpression shown in Figure 1F, while changes in the shed length *L_shed_* are negligible, which is also similar to *L_shed_* described in the *rheb*-overexpression experiment in Figure 1G.

In addition to testing just variations in the disc addition rate µ_g,_ we also looked to see if there were any possible synergistic effects between the parameters associated with shedding (the shedding rate α_s_ and the disposal rate µ_s_). In particular, we explored how variations of µ_g_ together with µ_s_, and then µ_g_ together with µ_s_ and α_s_, altered the growth and shed lengths at 14 days. Figure 4E illustrates that the simultaneous increase of both disc addition rate µ_g_ and disposal rate µ_s_ lead to an almost identical result as just increasing µ_g_ alone, which suggests that the addition of the disposal rate had little to no effect on the resulting growth and shed lengths. Figure 4F illustrates that increasing all three parameters together results in smaller growth and shed lengths, which is inconsistent with the experimental observations. Overall, results lead us to suggest that the disc addition rate, µ_g_, is the key parameter altered by Rheb-overexpression, suggesting that the mitochondrial hypothesis could explain ROS elongation when Rheb is overexpressed.

## Discussion

The mechanisms that individual cells use to calculate and maintain organelle and cell size remain poorly defined. Control of ROS size provides a tractable example of homeostatic size control that is uncomplicated by cell cycle influences, presents a conspicuous structure, and tools for quantitative analyses and genetic manipulation are available. Further, ROS renewal dynamics provides a powerful example of how mathematical modeling can be applied to cellular systems to provide insight into the mechanisms involved in cell size control. Defining the mechanisms that regulate ROS renewal and size is also critical for understanding how this process is disrupted in retinal degeneration diseases, where progressive outer segment shortening and loss drives photoreceptor degeneration. Extensions of our mathematical modeling framework, together with experiments aimed at uncovering mechanisms of ROS renewal, will aid in the formulation of hypotheses for how such diseases progress, leading to a description of valuable translational targets for treatment to slow or prevent vision loss.

## Materials and methods

### Animal Experimentation

This study was carried out in strict accordance with the recommendations in the Guide for the Care and Use of Laboratory Animals of the National Institutes of Health; the protocol was approved by the University of Massachusetts Amherst Institutional Animal Care and Use Committee. All fish lines were maintained in a mixed background of *AB*; *albino^b4/b4^* [33, 34]. We induced membrane bound mCherry expression in adult fish by a 45 minute incubation in 38.5°C water, after which the fish were returned to 28°C fish water that included 5 μg/ml Doxycycline (DOX), fish were exposed to DOX 20h/day in non-circulating fish system water with an air bubbler for 14 days, fish were fed each day during the four hours of recirculating water.

The rod-specific TetOn driver *Tg(Xla.rho:rtTA;TRE:GFP)* line and the *Tg(hsp70:HA-mCherry^TM^)* line have been described previously [10] [11]. The TRE:rheb^αBtx^ transgene construct was made by amplifying zebrafish *rheb* from first stranded total RNA from 30-day larvae, and tagged at the N-terminus with the alpha bungarotoxin αBtx-binding peptide WRYYESSLEPYPD. The p5E_TRE plasmid (Tet-On toolkit, [11]) was combined with pME Rheb^lllBtx^ into pTol Gateway destination vectors [35].

The *Tg(TRE:rheb*^αBtx^) germ-line transgenic line was created using the pTol system [36] by co-injecting the TRE:Rheb^αBtx^ pTol plasmid with *transposase* RNA into one-cell *alb^-/-^* embryos, germ-line founders identified and stable, germline, a single transgene copy transgenic line that exhibited minimal positional effect variegation was propagated.

Genotyping: Animals used are *Tg(Xla.rho:rtTA;TRE:GFP); Tg(hsp70:HA-mCherry^TM^)* without and with *Tg(TRE:rheb^aBtx^)*. Larvae fish (7 dpf) are genotyped for *Tg(Xla.rho:rtTA;TRE:GFP)* by GFP fluorescence following 4-7 dpf DOX treatment, and *Tg(hsp70:HA-mCherry^TM^)* by red lens fluorescence (*hsp70* is constitutively expressed in lens). Adult fish are genotyped as *Tg(TRE:rheb^aBtx^)* by PCR of DNA isolated from fin-clips.

### Tissue and immunohistochemistry

Eyes were removed from euthanized adult fish and fixed for 1.5 hour (h) in 3% paraformaldehyde. Eyes were embedded in 1.5% agar/5% sucrose and equilibrated in 30% sucrose. Samples were sectioned (thickness = 25) μm using a Leica cryostat. Sections were rehydrated with phosphate buffered saline (PBS), permeabilized for 4-6 h with 10% goat serum in PBS containing 0.1% Triton X-100 (Sigma-Aldrich, St. Louis, MO, USA), and incubated overnight at 4°C in primary antibody in PBS containing 0.1% Tween 20 (PBS-Tw; Sigma-Aldrich, St. Louis, MO, USA). Sections were washed the next day with PBS-Tw, incubated for at least 5 h with the appropriate secondary antibody in PBS-Tw, and washed again with PBS-Tw. Additional primary and secondary antibody combinations were incubated on the sections over subsequent nights and days. Following incubation of all necessary antibodies, samples were mounted with ProLong^®^ Gold Antifade Mountant (ThermoFisher Scientific, Waltham, MA, USA). Antibodies used: rabbit anti-GFP primary antibody at 1:2000 (ThermoFisher Scientific, Waltham, MA, USA) with corresponding Alexa Fluor^®^ 488-conjugated goat anti-rabbit secondary antibody at 1:2000 (ThermoFisher Scientific, Waltham, MA, USA), mouse IgG_1_ monoclonal anti-HA primary antibody at 1:1000 (Covance, Princeton, NJ, USA) with corresponding Rhodamine Red-conjugated goat anti-mouse IgG_1_ secondary antibody at 1:100 (Jackson ImmunoResearch Laboratories, West Grove, PA, USA) or Alexa Fluor^®^ 546 goat anti-mouse (Jackson ImmunoResearch Laboratories, West Grove, PA, USA), rabbit anti-RFP (Rockland, Limerick, PA, USA) with Alexa Fluor^®^ 546 goat anti-rabbit, mouse anti-acetylated tubulin (Sigma Aldrich) with Alexa Fluor^®^ 546 goat anti-mouse, and R6-5 mouse IgG_2A_ monoclonal anti-Rhodopsin primary antibody [37] at 1:200 with corresponding Alexa Fluor^®^ 647-conjugated goat anti-mouse IgG_2A_ secondary antibody at 1:100 (Jackson ImmunoResearch Laboratories, West Grove, PA, USA). Alexa Fluor^®^ 647-α bungarotoxin (ThermoFisher Scientific, Waltham, MA, USA) was used at a 1:150 dilution. DAPI (ThermoFisher Scientific, Waltham, MA, USA) was used at 0.5 μm/ml.

### Microscope image acquisition and analyses

Images were generated as z-stacks of optical sections using a Nikon A1R-SIMe confocal microscope with a 20x/0.75 NA or 40x/1.4 NA oil objective and processed with Nikon Elements software. Representative images are projections of a subset of the z-sections as described in figure legends. Measurements were acquired using Nikon Elements software. Measurements (growth, shedding, and axoneme length) were taken from at least 4 retinas (similar region) from individual fish. Growth measurements were collected from an eye from 5 *wts* (individual measurements: 44, 30, 35, 29, 30) and 7 *TRE:rheb* transgenics (individual measurements: 45, 34, 39, 38, 25, 42, 49). Shedding measurements were collected from a similar region of retina from 5 *wts* (individual measurements: 57, 36, 40, 36, 46) and 7 *TRE:rheb* transgenics (individual measurements: 49, 25, 36, 43, 22, 38, 39). Axoneme length was made from a similar region of retina from 4 *wts* (number of axonemes measured per individual: 72, 59, 56, 33) and 4 *TRE:rheb* transgenics (number of axonemes measured per individual: 89, 30, 72, 16).

### Statistical analysis

Prism 8 software was used for statistical analyses (Welch’s *t*-test) and generating graphs.

### Model output (conversion between discs and length)

Since our ODE model describes discs over time, and we want to express our results in terms of length and provide a simple calculation for converting between disc number and length. In particular, we use measurements of disc-disc spacing and disc thickness for murine ROS, studied using cryoelectron tomography [32]. Scaling disc number to total length is given by

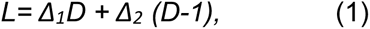

where *L* is length, *D* is the number of discs, and *Δ_1_* = 22.5 nm and *Δ_2_* = 10.0 nm represent the average thickness of a disc and the disc-disc spacing, respectively [38]. For our model, the total length of the rod, *L_tot_*, at any time *t* is a function of the total number of discs in both the G and M compartments at time *t* such that

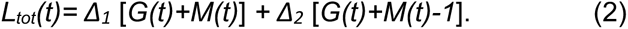

We initialize our model with discs in the *M* and *G* compartments such that *M(0)* > 0, *G(0)* > 0, and the initial length L_int_ *= Δ_1_* [*G(0)+M(0)*] *+ Δ_2_* [*G(0)+M(0)-1*], where *L_int_* corresponds to the length of the rod at the time of heat shock. Then, at any time *t*, we can determine the shed length *L_shed_*(*t*) as

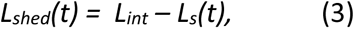

where *L_s_(t)* corresponds to the length associated with discs in the recycling compartment *S* time *t*. The growth length *L_growth_(t)* at time *t* is determined by subtracting the shed length from the total length such that

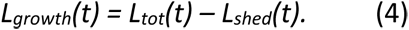

## Abbreviations

αBtx: alpha bungarotoxin
DOX: doxycycline
ODE: ordinary differential equation
ROS: rod outer segment
RPE: retinal pigmented epithelium
SD: standard deviation

## Acknowledgements

This research was supported by a collaborative research grant from the National Science Foundation (NSF), Division of Biological and Biomedical Sciences (Award ID 1951420).

