## Supplemental Data for "Overexpression of Rheb, a positive Tor regulator, reveals principles of rod outer segment size control"

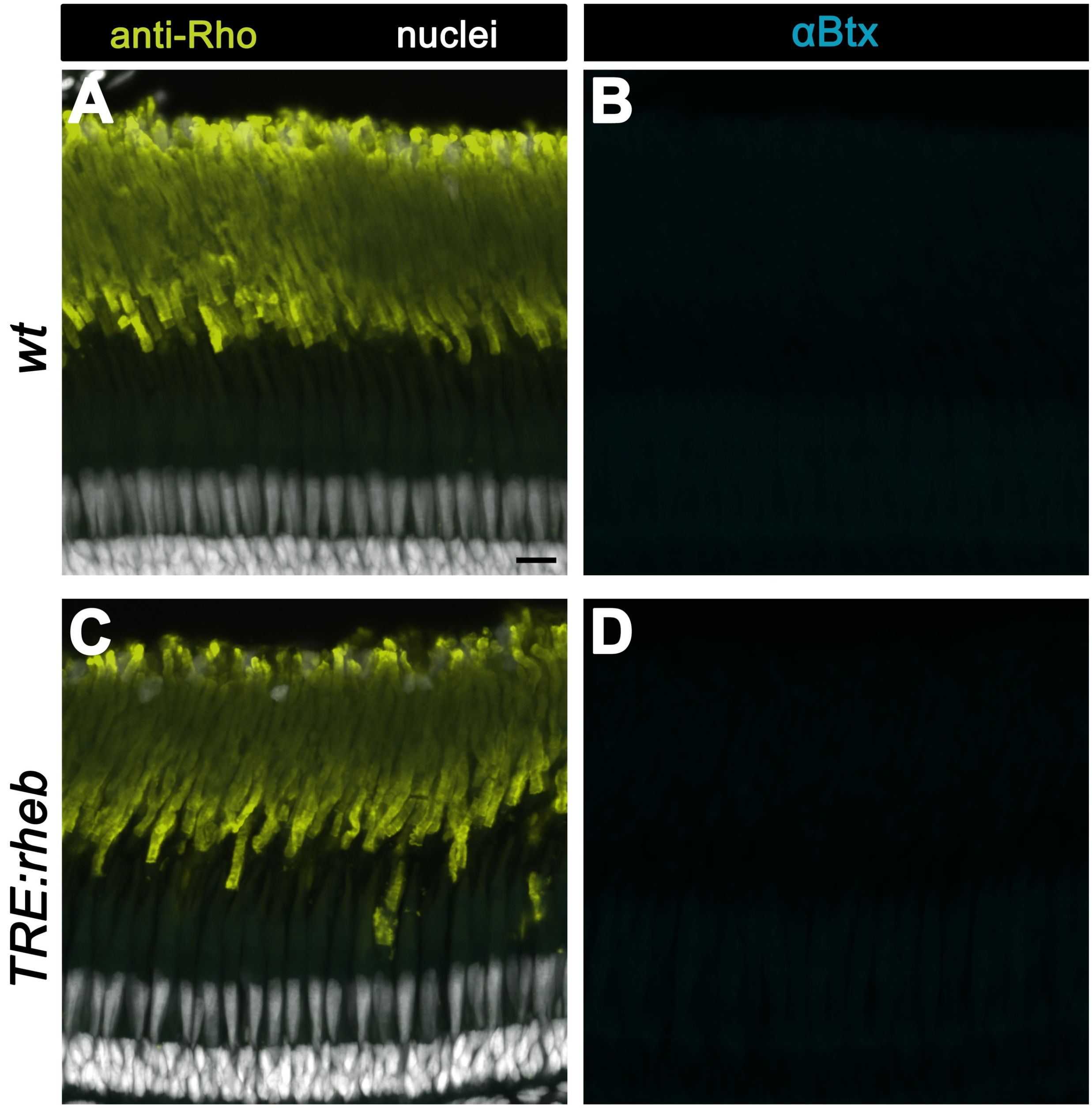


**Supplemental Figure 1**: Confocal z-projections (z = 4 μm) of untreated *wild-type* (*wt*) and *TRE:rheb* transgenic retinas labeled with α–bungarotoxin (αBtx), the ROS is labeled with anti-Rhodopsin (anti-Rho) antibodies, nuclei labeled with DAPI. Scale bar, 10 μm.

**Supplemental material**

The system of ordinary differential equations (ODEs) used to describe the dynamics of the ROS is given by

$$\frac{dG}{dt} = \mu_g(G, M, S) - \alpha_g G$$

$$\frac{dM}{dt} = \alpha_g G - \alpha_s(G, M, S)M$$

$$\frac{dS}{dt} = \alpha_s(G, M, S)M - \mu_s S$$

Each equation describes the rate of change of discs in the G, M, and S compartments, respectively. In general, we assume that the disc addition rate $\mu_g$ and the shedding rate $\alpha_s$ may be dependent on the number of discs in the growing compartment G, the mature compartment M, and the shed compartment S. For simplicity in the current study, we have assumed that all model parameters are constants. By treating the model in this way, the goal is to describe qualitatively (and perhaps even quantitatively), how ROS baseline dynamics (defined below) are altered for changes in model parameters.

**Determination of the baseline parameter set:** To calibrate our model, such that starting from any initial condition results in the growth and shed lengths found in wild type (*wt*) experimental results at 14 days (see Fig 1F and 1G, respectively, in the main text), we fixed $\alpha_g$ = 0.85 μm/day (which corresponds to the average growth length over 14 days from Fig 1F), while performing a parameter sweep over the other three model parameters along the ranges listed below. After recording all parameter sets which gave us the growth and shed lengths within experimental range, we took the averages for each parameters $\mu_g$, $\mu_s$ $\alpha_s$ and used these as our baseline parameter set.

**Parameter set ranges for parameter sweep and the resulting baseline values:**

Range for $\mu_g$ was 30 to 180 discs per day

Range for $\alpha_s$ was 0.1 to 1.4 μm/day

Range for $\mu_s$ was 0.1 to 1.4 μm/day

Baseline set: $\alpha_g$ = 0.85 μm/day $\mu_g$ = 82 discs/day, $\mu_s$ = 0.96 μm/day, and $\alpha_s$ = 0.17 μm/day

**Determining model equilibrium/set length:** Setting the time derivatives in the above equations equal to zero corresponds to the model's "equilibrium" state, such that G, M, and S do not change in time. The resulting 3-D system of equations with 3 unknowns G, M, and S, can be solved to give (G*, M*, S*) = ($\mu_g/\alpha_g$ , $\mu_g/\alpha_s$, $\mu_g/\mu_s$) , where "*" indicates the state of each variable at equilibrium. After standard analysis of the equilibrium's stability (determination of the signs of the eigenvalues of the Jacobian Matrix at this equilibrium [25], we determine that the equilibrium is globally asymptotically stable (i.e., the eigenvalues are negative), such that for any fixed parameter set and starting from any initial conditions (initial disc number in each compartment G, M, and S), the same equilibrium/set length will be reached.

From the equilibrium expression, perturbations (and, in particular, increases) in the maturation rate $\alpha_g$ only impacts developing discs in the G compartment by decreasing the number of new discs added at the base, and results in no change in the number of discs in the mature compartment M. The overall result is a small decrease in the total length of the ROS at equilibrium; thus, we exclude this simulation test in the paper.
